# KIR*BLOOM: Accurate KIR genotyping using a new copy number-aware integrated genotype likelihood framework

**DOI:** 10.64898/2026.04.15.718735

**Authors:** Yomna Gohar, Alberto Descalzo Garcia, Katherine M. Kichula, Paul J. Norman, Alexander Dilthey

## Abstract

Killer-cell immunoglobulin-like receptor (KIR) genes, key modulators of natural killer (NK) cell activity, play critical roles in immune response and disease susceptibility. Accurate KIR genotyping from short-read sequencing data remains challenging because of high sequence similarity among genes, extensive copy number variation, and substantial allelic diversity. Here, we present KIR*BLOOM, a likelihood-based approach for KIR genotyping from short-read data that models read depth and sequencing error across alternative genotype configurations. KIR*BLOOM first identifies KIR-relevant read pairs, maps them to a KIR allele database, and reduces the candidate allele space by excluding alleles unlikely to be present. It then infers gene copy number and selects alleles under the inferred copy-number constraints. Finally, variant calling is used to refine CDS sequences and identify potential novel alleles. We evaluated performance on 45 whole-genome sequencing samples with haplotype-resolved assemblies from the HPRC or HGSVC, using Immuannot-derived annotations as ground truth. KIR*BLOOM achieved 99.85% precision, 99.92% recall, and a Jaccard index of 99.77% for copy-number inference. At five-digit allele resolution, it achieved 92.73% precision, 92.69% recall, and an 87.29% Jaccard index, outperforming T1K, GraphKIR, and Geny. Together, these results demonstrate that KIR*BLOOM enables highly accurate KIR genotyping from short-read sequencing data.

## Introduction

Killer-cell immunoglobulin-like receptors (KIRs) are a family of receptors expressed on natural killer (NK) cells and a subset of T cells (Campbell and Purdy 2011) that interact primarily with major histocompatibility complex (MHC) class I molecules on host cells (Vilches and Parham 2002). These interactions are central to immune surveillance, enabling NK cells to balance activation and inhibition, thereby detecting and eliminating cells with intracellular abnormalities such as viral infection or malignant transformation (Benson and Caligiuri 2014; Trowsdale et al. 2015). KIR receptors are encoded by KIR genes located on chromosome 19q13.4. Since their discovery in the early 1990s, 17 KIR genes have been identified. They include eight inhibitory receptors (KIR2DL1, KIR2DL2, KIR2DL3, KIR2DL5A, KIR2DL5B, KIR3DL1, KIR3DL2, and KIR3DL3), seven activating receptors (KIR2DS1, KIR2DS2, KIR2DS3, KIR2DS4, KIR2DS5, KIR3DS1, and the dual-function KIR2DL4), and two pseudogenes (KIR2DP1 and KIR3DP1) (Béziat et al. 2017). Individuals can carry diverse combinations of 8 to 17 KIR genes. In addition, KIR genes exhibit extensive allelic polymorphism and gene copy number variation, ranging from complete absence to up to four copies of a single gene per individual, resulting in hundreds of observed haplotype structures. Despite this diversity, KIR haplotypes are broadly classified into group A and group B, which differ in gene content and functional balance. Group A haplotypes are relatively conserved and enriched for inhibitory receptors, typically containing a single activating gene (KIR2DS4), whereas group B haplotypes are more diverse and harbor multiple activating KIR genes, contributing to greater variability in immune responses (Hollenbach et al. 2012; Hsu et al. 2002). Genetic variation within the KIR locus has been associated with a wide range of clinically relevant phenotypes, including susceptibility to infectious diseases (Khakoo et al. 2004; Martin et al. 2002), autoimmune disorders (Hollenbach et al. 2009), and cancer (Purdy and Campbell 2009; Liu et al. 2015). In pregnancy, KIR receptors play a critical role in regulating uterine NK cell interactions with fetal trophoblasts, thereby influencing placental development and the risk of disorders such as pre-eclampsia (Kulkarni et al. 2008).

Genotyping KIR genes remains particularly challenging because the KIR locus combines extensive allelic polymorphism, high sequence homology between genes, and substantial structural variation (Roe et al. 2020). While a central database of KIR alleles exists (IPD-KIR) (Robinson et al. 2019) currently containing 2,286 alleles, However, even with this reference, precise genotype inference remains difficult because many alleles are distinguished by only small sequence differences, including synonymous substitutions within coding regions or variants confined to intronic or other non-coding regions. Closely related genes such as KIR3DL1/KIR3DS1 and KIR2DL5A/KIR2DL5B (Song et al. 2023), for example, share considerable sequence similarity, increasing the risk of ambiguous read assignment. In addition, about 40% of the alleles in the database are incomplete. Together, these factors, including the extensive copy number variation of KIR genes described above, underscore the need for specialized and accurate genotyping approaches.

Among the currently available tools for allele-level KIR genotyping from short-read sequencing data, PING (Marin et al. 2021) and T1K (Song et al. 2023) are among the best established, with additional newly developed methods such as Geny (Zhou et al. 2024a) and GraphKIR (Lin et al. 2023) also contributing to the available set of approaches. PING is a KIR-specific workflow that extracts KIR reads and then estimates gene copy number and allele genotypes, whereas T1K is developed for both HLA and KIR alleles inference by estimating allele abundances from read alignments and supports RNA-seq, whole-genome, and whole-exome sequencing data. However, both approaches retain important limitations. PING has been reported to require substantial manual parameter tuning and to be sensitive to cohort-dependent thresholds and copy-number normalization assumptions. T1K is efficient and broadly applicable, but recent evaluations indicate that complex KIR copy-number states are not modeled robustly, which can lead to undercalling of gene copies, particularly when copy number exceeds two. In addition, newer tools such as Geny and GraphKIR have not yet been extensively benchmarked across diverse real-world short-read datasets. Together, these limitations restrict accurate fine-mapping of KIR-associated phenotypes to their underlying genotypes. Consequently, there remains a clear need for improved tools for accurate high-resolution KIR genotyping.

Motivated by this need, we developed KIR*BLOOM (bounded likelihood optimization for observed mixture), a method that explicitly accounts for ambiguously mapping reads and first infers KIR gene copy number and then uses this information to constrain a likelihood function for identifying the combination of KIR alleles that best explains the observed read mixture. KIR*BLOOM was designed to be cohort-agnostic and therefore does not require manual parameter tuning or predefined thresholds. Moreover, it focuses on recovering the correct coding sequence (CDS) of KIR alleles and further refines allele sequences through variant calling.

## Results

### Development of high accuracy KIR genotyping method

We developed KIR*BLOOM, a method for inferring KIR gene copy numbers and alleles from short read sequencing data. We formulate KIR genotyping as an optimization problem over copy-number vectors over alleles of multiple genes; For possible copy-number configurations across candidate alleles, we seek the configuration that best explains the observed read alignments. For any proposed copy-number vector, KIR*BLOOM constructs, for each allele, a coverage profile and error profile (reflecting mismatches, Insertion, deletions and clipping in aligned reads) by probabilistically assigning reads to alleles, thereby accounting for ambiguous mappings in the highly homologous KIR locus. The resulting allele-specific coverage and error profiles are then scored by comparing them to a background expectation estimated from a stable reference region on chromosome 17 (Prodanov et al. 2025), and statistical optimization is carried out to find the copy number vector associated with the best overall coverage and error profile (Figure 1).

**Figure 1.**
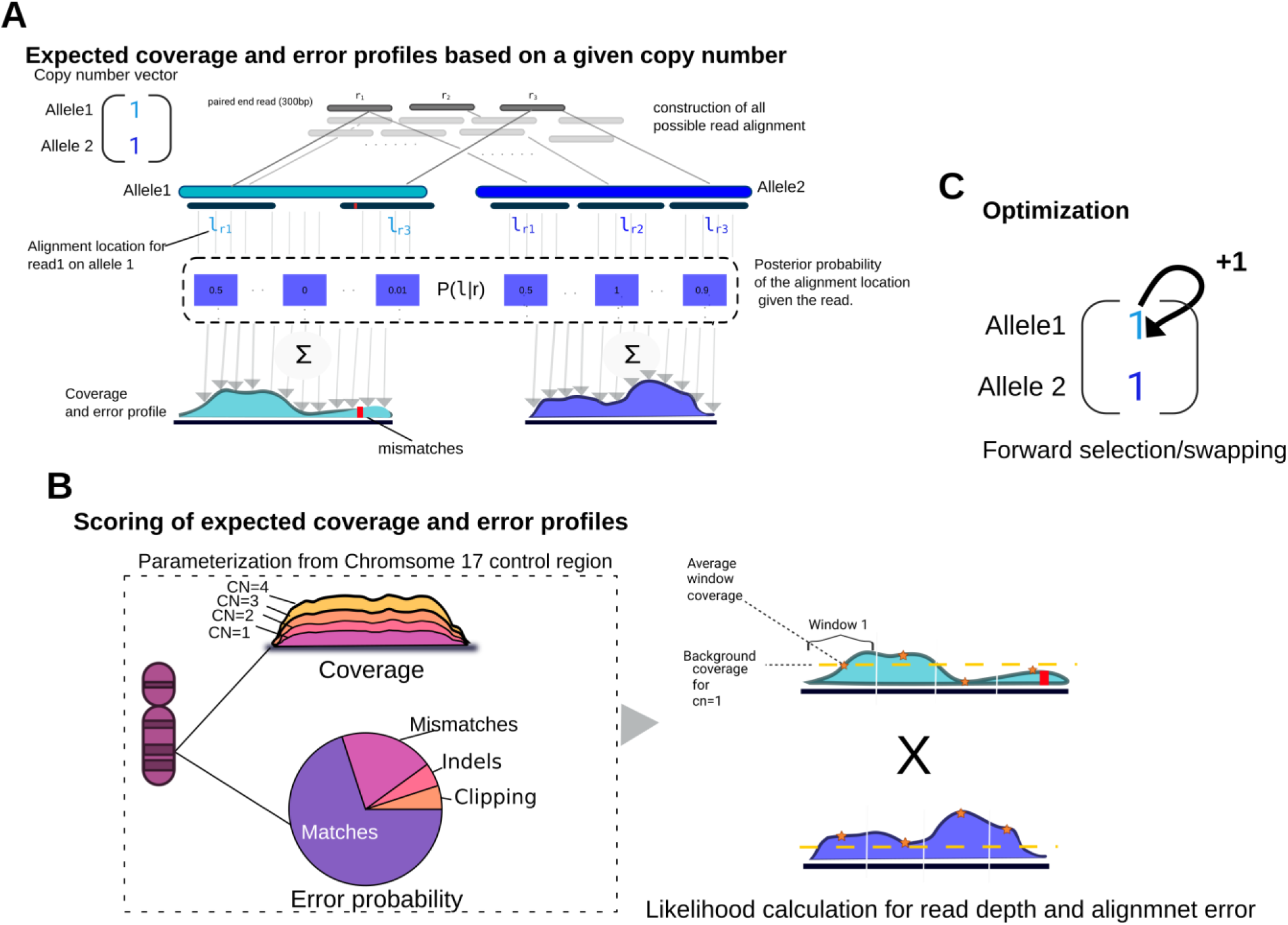
Overview of the three main steps of the KIR*BLOOM algorithm. **(A)** For a proposed allele copy-number configuration, KIR*BLOOM probabilistically assigns ambiguously mapping reads across candidate alleles and constructs allele-specific expected coverage and alignment-error profiles. **(B)** These profiles are scored against background expectations estimated from a stable control region on chromosome 17, incorporating both read depth and alignment-error patterns, including mismatches, insertions, deletions, and clipping. **(C)** KIR*BLOOM then optimizes the copy-number configuration by forward selection and swapping to identify the solution that best explains the observed read alignments. Together, these three steps form the core inference framework used for copy-number and downstream allele genotyping.

KIR*BLOOM begins by extracting KIR-candidate read pairs from WGS. Then it generates all possible read alignment to a KIR*BLOOM allele database from the KIR*BLOOM reference. The KIR*BLOOM reference combines (i) a KIR*BLOOM allele database which is derived from IPD-KIR, completing CDS-only alleles by inferring missing segments from the closest related alleles, and further expanded with additional alleles extracted from T2T and GRCh38 and (ii) the T2T and GRCh38 human genomes in which the KIR locus is masked, thereby discouraging spurious mappings to KIR alleles (Methods, Supplementary Notes). The resulting alignments are processed through three stages: (1) allele filtering to reduce the solution space and runtime, (2) copy-number estimation using a likelihood model that selects the gene copy-number vector that yields a read-depth profile most consistent with the expectation, and (3) exon-level (five-digit) allele inference, in which the inferred copy numbers impose explicit bounds on the downstream allele-resolution step; during exon-level inference, the same likelihood framework already used for copy number estimation is used together with the error profile rather than depth alone (Supplementary Figure 1).

### Performance on gene copy number

We ran KIR*BLOOM on short-read whole-genome sequencing data from 97 samples with available haplotype-resolved assemblies from the HPRC and HGSVC resources (Supplementary Table 1). Of these, 52 samples were used for method development and 45 for independent evaluation; all presented evaluation results are based on the 45 independent samples. The assemblies (Supplementary Table 1) were used to derive ground-truth genotypes with Immuannot (Zhou et al. 2024b). For comparison, we evaluated three currently available methods, T1K, Geny, and GraphKIR, on the 45 independent evaluation samples. For each method, we assessed agreement between the predicted and reference gene copy numbers (CN) using absolute CN accuracy, precision, recall, and the Jaccard index (Methods). Absolute CN accuracy was defined, for each gene, as the proportion of samples in which the predicted copy number exactly matched the ground truth for each gene (i.e., for each gene, an exact-match indicator of 1 for a match and 0 otherwise, summed over all samples and divided by the total number of samples). Metrics were computed separately for each gene, except for KIR2DL5A and KIR2DL5B, whose copy numbers were aggregated and evaluated jointly as KIR2DL5.

KIR*BLOOM and T1K achieved 100% gene detection accuracy, correctly identifying the presence or absence of every gene in the reference set across all evaluated samples. GraphKIR also achieved 100% accuracy for all genes except KIR2DS1, for which accuracy dropped to 84.4%. Geny showed the lowest overall gene detection performance, with an average accuracy of 98.3% and a minimum per-gene accuracy of 93% for KIR2DS1 (Supplementary Table 2). For exact copy number prediction for each gene, KIR*BLOOM showed very high copy-number prediction performance, achieving an overall absolute accuracy of 99.58%, compared with 93.61% for T1K, 93.4% for Geny, and 97.50% for GraphKIR (Figure 2; Supplementary Figure 2). Across 867 gene copy-number calls, KIR*BLOOM made only 2 errors (2 false positive) and missed one call (1 false negative), which we investigated in detail. One error involved overestimation of the copy number of KIR2DS4 by one. This occurs due to a KIR3DL1–KIR2DS4 fusion gene (Hung et al. 2024) which is absent from the reference dataset (IPD-KIR) and couldn’t be detected by immuannot. For the other two errors, there was no clear reason. We further assessed whether excluding novel alleles derived from the GRCh38 and T2T assemblies affected the results; however, the results remained identical.

**Figure 2.**
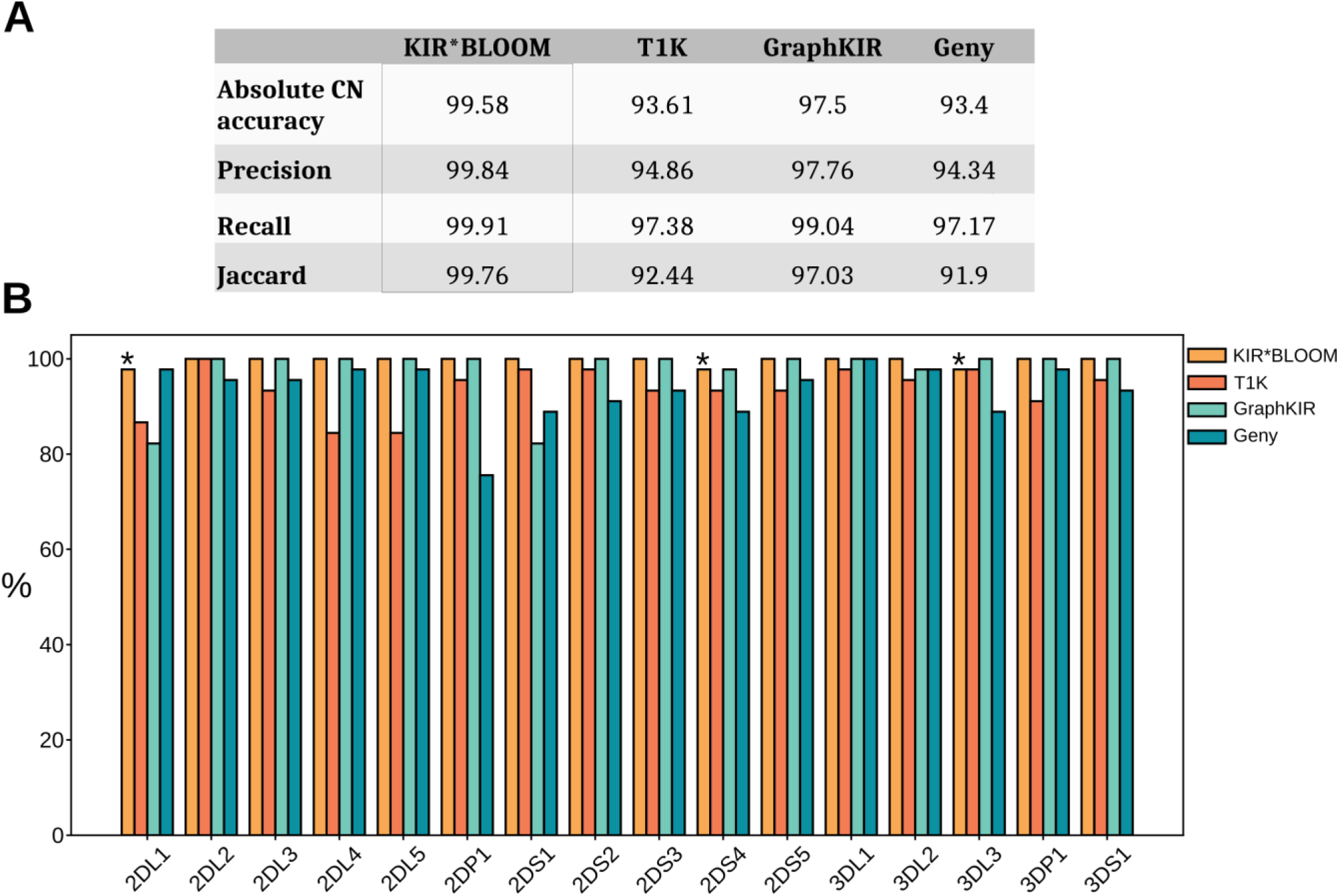
Evaluation of gene copy-number inference performance. **(A)** Summary of overall copy-number performance for KIR*BLOOM, T1K, GraphKIR, and Geny. Performance metrics were computed separately for each gene and then averaged across genes. **(B)** Gene-stratified absolute copy-number accuracy for each method, shown as the percentage of exact copy-number matches relative to the ground truth. An asterisk indicates genes for which KIR*BLOOM did not achieve 100% performance.

### Performance on five-digit allele inference

We next evaluated performance for five-digit allele inference, which reflects prediction of the correct coding-sequence (CDS). We assessed our method using allele match, recall, precision, and the Jaccard index (Methods). Allele match was defined per sample and per gene as 1 if all alleles were predicted correctly with their corresponding copy numbers, and 0 otherwise. KIR*BLOOM achieved the best average five-digit typing performance using the KIR*BlOOM allele database ( Methods). For allele match specifically, KIR*BLOOM achieved 90.95%, notably exceeding T1K and Geny, which achieved 80.71% and 73.47, respectively, while GraphKIR showed the lowest performance at 62.52% (Table 1).

**Table 1:** Comparison of KIR*BLOOM, T1k, GraphKIR and Geny on 45 evaluation samples.

|  | KIR*BLOOM |  | T1K |  | GraphKIR |  | Geny |  |
| --- | --- | --- | --- | --- | --- | --- | --- | --- |
|  | Before VC | After VC | Before VC | After VC | Before VC | After VC | Before VC | After VC |
| Allele match | 88.09 | 90.95 | 78.16 | 80.71 | 62.52 | - | 73.47 | - |
| Jaccard | 83.06 | 87.29 | 77.8 | 80.76 | 57.33 | - | 72.33 | - |
| Precision | 90.05 | 92.73 | 85.53 | 87.31 | 70.33 | - | 82.42 | - |
| Recall | 89.98 | 92.69 | 89.09 | 90.89 | 70.8 | - | 84.59 | - |

Compared with T1K after variant calling and correction, KIR*BLOOM achieved higher precision (13 genes), allele match (16 genes), recall (9 genes), and jaccard index (15 genes) (Figure 3). T1K showed a higher jaccard index for two genes, KIR3DP1 and KIR2DL2, by 2.7% and 7%, respectively. However, although T1K performed better for KIR3DP1, this gene, together with KIR3DL3, had the highest number of errors overall for all evaluated methods. Compared with Geny, KIR*BLOOM outperforms it in 15 genes for precision, 14 genes for recall, 15 genes for allele match, and 15 genes for the Jaccard index. For the Jaccard index, Geny outperformed KIR*BLOOM for the genes KIR2DL4 and KIR2DL5A by 1% and 2.3%, respectively. For genes KIR2DL3, KIR2DS1, KIR2DS3, KIR3DS1, and KIR3DL1, KIR*BLOOM achieved perfect performance (100%) across all evaluated metrics. In contrast, T1K achieved perfect performance across all evaluated metrics only for KIR2DL2. For this gene, KIR*BLOOM made a single error involving a novel allele, where it failed to identify the correct variant. GraphKIR achieved perfect performance for KIR3DS1, like KIR*BLOOM, whereas Geny did not achieve perfect performance for any gene.

**Figure 3.**
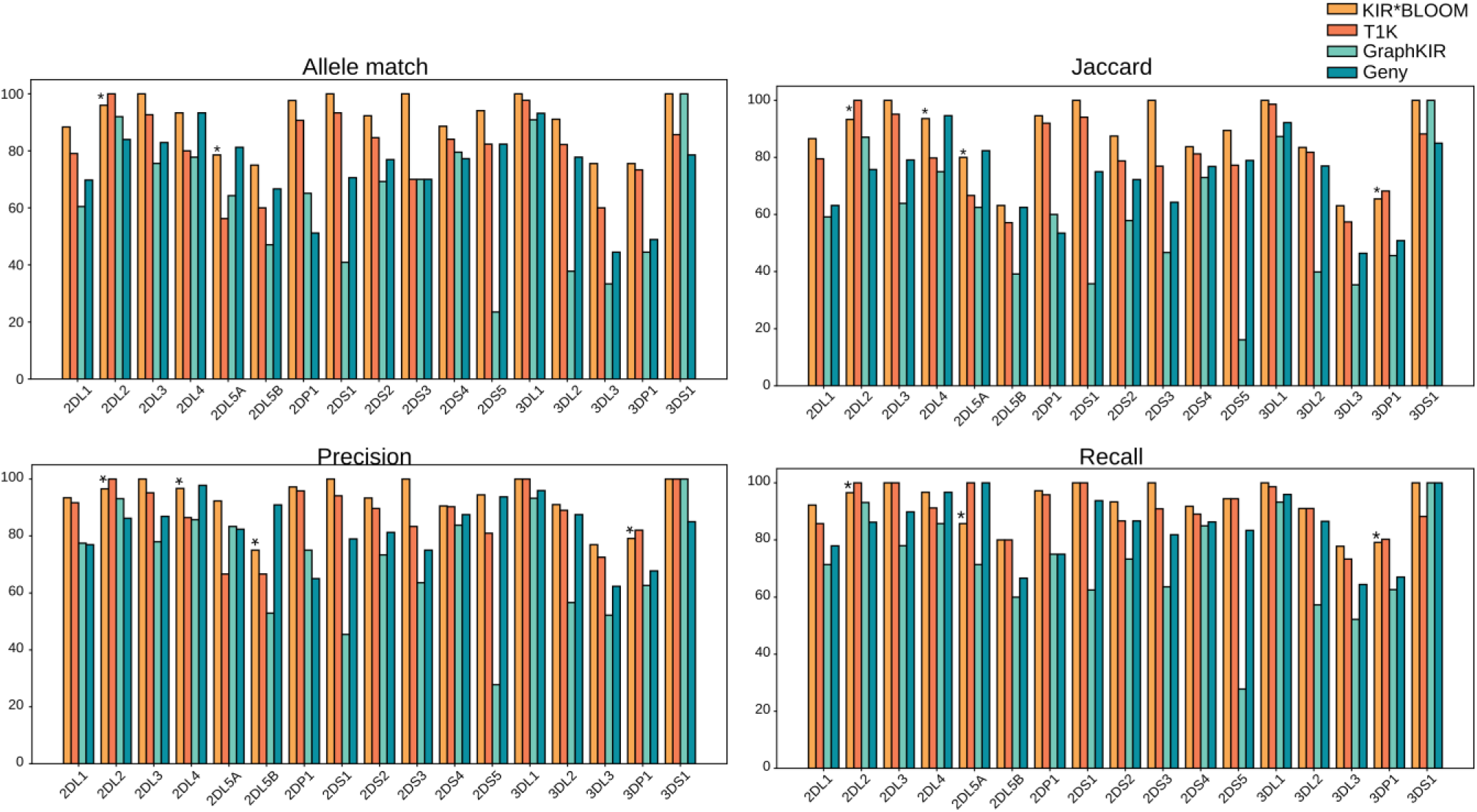
Gene-wise performance of KIR inference at the 5-digit level. An asterisk indicates genes for which KIR*BLOOM was outperformed by at least one other method.

We then evaluated the performance of the different algorithms when using only an IPD-KIR-based database (i.e., not including the alleles in the extended KIR*BLOOM allele database) and without variant calling; here, KIR*BLOOM outperforms all tools on all metrics, with the exception of recall, where T1K achieves the best overall result (90.89% compared to 88.81% for KIR*BLOOM) (Supplementary Table 3). Still based on the IPD-KIR database, we then evaluated the effect of variant calling, which is only supported by KIR*BLOOM and T1K. In this comparison, KIR*BLOOM outperformed T1K on all metrics including recall. Finally, comparing KIR*BLOOM using an IPD-only database to its full extended database showed an average performance increase of 1% across all metrics. Overall, the inclusion of additional alleles increases true positives by eight. The additional alleles contributed most notably to improved performance for KIR3DL3. This improvement was primarily driven by the addition of a complete KIR3DL3*005 to the KIR*BLOOM allele database; in-depth investigation on the development set (which was not used in any validation experiments) showed that the IPD-KIR version of this allele lacked intron 1. We therefore added a corrected version that included intron 1. This increased the number of true positives by five. Another added allele that contributed to the improvement was KIR2DL5B*00601, which contained a novel intronic variation compared with the IPD-KIR version. Together, these results indicate that the improved performance of KIR*BLOOM reflects an improved performance of the underlying genotyping algorithm as well as that of an improved database.

Next, we characterized the effect of the variant calling step, which may, in principle, (i) enable the detection of novel alleles not present in the database; (ii) correct errors made by allele selection step; (iii) introduce false-positive variants. Across the analyzed samples, 39 alleles carried a novel coding sequence (CDS) (Supplementary Table 4). Of these, KIR*BLOOM method correctly recovered 10, whereas T1K recovered 17. Although T1K recovered more novel CDS sequences than our method, both approaches resolved fewer than half of all novel CDS sequences. For KIR*BLOOM, recovery was particularly difficult for genes with copy numbers greater than one were the method only reports unambiguous allele call (Methods). Overall, variant calling reduced the number of false negatives for both methods. For KIR*BLOOM, the number of false negatives across all loci decreased from 89 to 73, corresponding to an approximately 18% reduction, and performance increased across all evaluation metrics by an average of 3%. For T1K, false negatives decreased from 109 to 92, corresponding to an approximately 16% reduction. However, For T1K, this improvement was partly offset by unnecessary sequence changes introduced into alleles whose CDS sequences had already been correctly assigned. For example, this could occur for homozygous alleles where sequence modifications were applied based on the estimated allele frequencies (Methods).

The IPD-KIR database includes alleles with (i) undefined intron sequences; (ii) incomplete exon sequences; or (iii) both. When constructing the KIR*BLOOM allele database, the missing regions were interpolated from closely related alleles (see Methods and Supplementary Notes). We investigated whether KIR*BLOOM could recover the alleles’ complete CDS in cases in which the evaluation samples carried these incomplete alleles, because these interpolated regions may not fully represent the true sequence, alignments to such alleles may be suboptimal. We therefore assessed whether the CDS of these incomplete alleles could still be identified correctly. Across all samples, 8 occurrences representing 5 alleles involved partially or completely missing exons cases. KIR*BLOOM correctly recovered the CDS sequence in all 8 instances, whereas T1K recovered 2 instances corresponding to KIR2DS4*00104. GraphKIR and Geny missed them all (considering the predicted allele name and not CDS). Among alleles with complete CDS sequence but missing intronic sequence, there were 74 instances across the samples. KIR*BLOOM recovered the correct CDS sequence in 60 instances, compared with 51 instances for T1K, 19 instances for GraphKIR and 30 instances for Geny (Supplementary Table 5).

### Analysis of KIR bloom error mode

KIR*BLOOM produced 74 false positives and 43 false negatives. In nearly all cases, each false negative was accompanied by a corresponding false positive, indicating that most errors arose from allele misassignment rather than complete omission. There were three exceptions: one resulted from underestimation of the KIR2DL1 copy number by one, and two involved switching between KIR2DL5A and KIR2DL5B alleles. In the latter two cases, a false positive was still generated for each false negative, but not within the same gene. In addition, two false positive calls arose from copy number overestimation: one for KIR3DL3 and one for KIR2DS4. The KIR2DS4 overcall is explained by a KIR3DL1–KIR2DS4 fusion that is absent from the IPD-KIR reference and therefore was not detected by Immuannot.

To understand the causes of the 73 false negatives produced by KIR*BLOOM, we classified them into four categories: loss of the true allele during solution-reduction filtering, failure to recover a novel coding sequence (CDS) by variant calling, ambiguous genotype configurations, and other gene-specific causes.

The most frequent error class was failure to rescue novel alleles by variant calling (Figure 4), which accounted for 39.7% (n = 29) of all false negatives. In KIR*BLOOM, a variant is called only when at least 75-85% of aligned reads support the sequence change (Methods). This stringent threshold can prevent recovery of novel CDS sequences when reads from the novel allele are distributed across multiple selected alleles, reducing the apparent support for the variant below the calling threshold. This problem was particularly pronounced in KIR3DP1, which had the highest number of novel CDS alleles (n = 9). In these cases, the presence of a novel allele often also led to incorrect selection of the second, non-novel allele. The second most common error class was ambiguous genotype configurations, which represented 32% of all false negatives (n=24). An ambiguous genotype occurs when two different pairs of alleles produce an identical diploid base-level sequence, while the heterozygous sites distinguishing the two configurations are separated by distances greater than the maximum fragment length, preventing read-based phasing (Supplementary Figure 3). Consequently, both allele pairs explain the data equally well and reads align to both without alignment errors on the exons. KIR*BLOOM therefore selects the allele pair with the highest likelihood, even when the improvement over the alternative combination is only marginal. This pattern was especially overrepresented in KIR3DL3, where it explained 18 of the 20 false negatives observed for this gene. It was also seen in KIR3DP1 (n=2) and KIR3DL2 (n=4), accounting for 10.5% and 50% of their false negatives, respectively.

**Figure 4.**
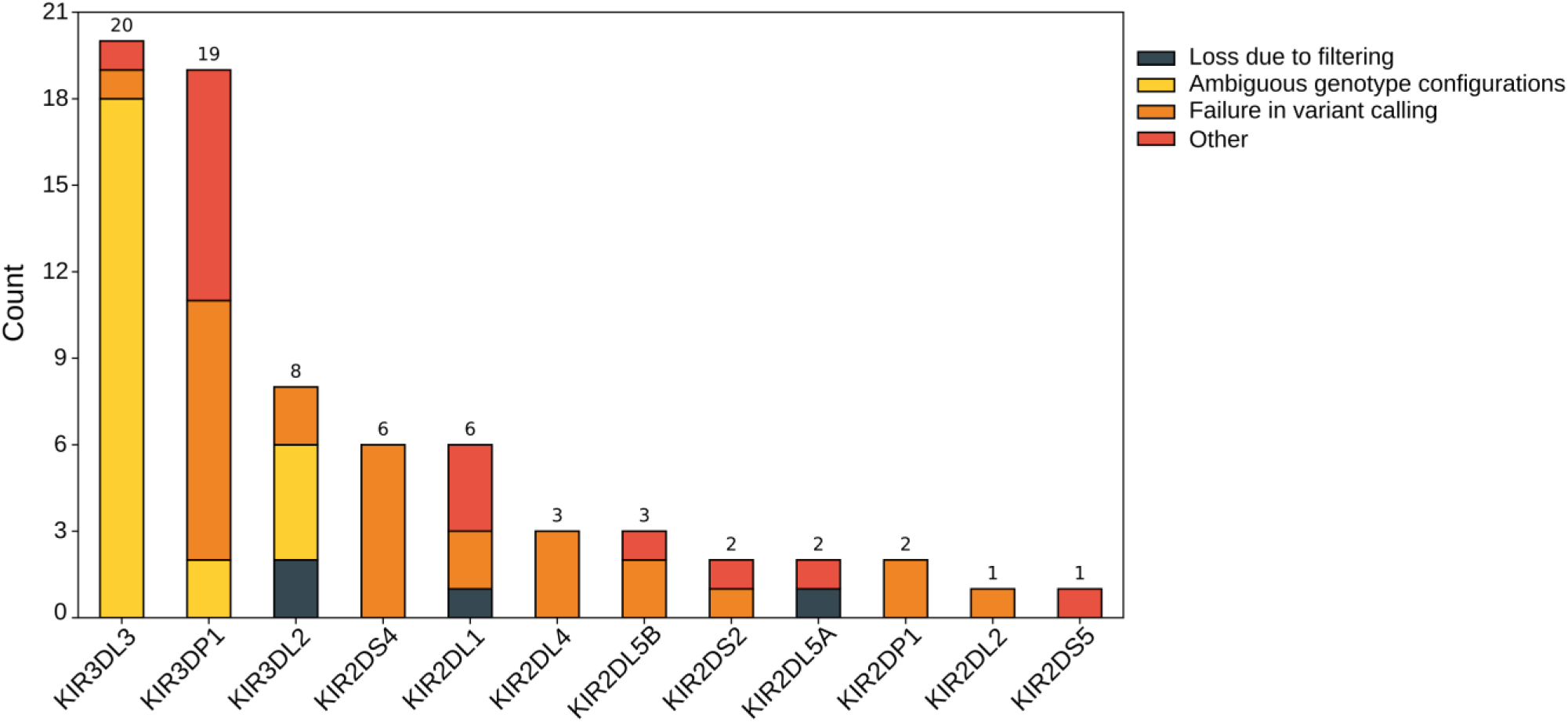
Gene-wise stratification of KIR*BLOOM false negatives by source.

The remaining false negatives were mainly caused by allele loss during filtering and another set of heterogeneous causes which mostly reflected limitations of the reference representation rather than a systematic weakness of the inference model. In several cases, incomplete reference sequences, particularly missing intronic regions in the IPD-KIR database, led to suboptimal read mapping and caused the correct allele either to be filtered out or not selected during inference. This was observed for KIR2DS2*005, KIR3DP1*004 (n=3), KIR2DL5A*01201, KIR3DL2*00601, and KIR3DL2*01004. Additional errors occurred when the presence of a novel allele led to loss of the second allele in the genotype. This was most frequent in KIR3DP1 (n=5) and was also observed in KIR3DL3 (n=1) and KIR2DL1 (n=1). The remaining false negatives arose from several sources. One was caused by copy-number misassignment during copy-number inference, as mentioned earlier. Another involved KIR2DL1, where true structural variation in exon composition made the genotype inherently ambiguous. Two additional cases reflected confusion between the highly similar paralogous genes KIR2DL5A and KIR2DL5B, with one misassignment in each direction. Finally, two false negatives, in KIR2DL1 and KIR2DS5, had no clear explanation.

## Discussion

Variations in KIR genes shapes NK-cell immunity. Accurate genotyping of the KIR locus remains technically challenging because the region combines extensive gene-content variation, high allelic polymorphism, copy-number variation, and strong sequence homology between related genes, and these challenges are further compounded by reference databases that do not yet fully represent the breadth of KIR allelic diversity. In this study, we developed KIR*BLOOM and evaluated its performance on whole-genome sequencing data from 97 real-world samples with available assemblies. We show that KIR*BLOOM addresses several of the major challenges of KIR genotyping by enabling accurate inference of gene presence and absence, showing an excellent performance for gene copy-number inference, and outperforming existing methods in CDS-level sequence prediction, particularly through the use of variant calling to recover correct CDS sequences. Nevertheless, two important limitations remain, both largely driven by incomplete reference representation: genotype ambiguity and the presence of novel alleles.

The strong performance of KIR*BLOOM was achieved in part through the combination of an extended reference database and variant calling. Interpolating regions missing from the IPD-KIR reference using closely related alleles in the extended database improved reference completeness. This enabled correct prediction in cases where the missing regions included CDS sequence, which was a factor limiting the performance of T1K, and also improved detection of alleles with missing sequence in non-coding regions by facilitating read mapping and allowing such alleles to remain under consideration during inference. Nevertheless, this approach does not fully resolve all cases, because when the inferred reference sequence remains substantially different from the true allele, alignment can still be suboptimal, leading to failure in correct allele prediction. Variant calling is an important extension that remains absent from many KIR genotyping methods, although it is implemented in T1K. Here, we applied stringent criteria for sequence correction to minimize false variant calls, particularly in genes with copy number greater than one, where reads from a novel allele may be distributed across multiple inferred alleles and generate misleading evidence for sequence changes. We therefore prioritized specificity and called variants only when supported by strong evidence. Although this conservative strategy reduced recovery of true novel variants to approximately 25%, it limited erroneous sequence corrections and preserved the overall reliability of allele inference.

The predominant source of error was genotype ambiguity, which had the greatest impact on KIR3DL3, KIR3DL2, and KIR3DP1. KIR3DL3 and KIR3DP1 also showed comparatively weak performance across other methods, suggesting that these cases reflect a broader difficulty of KIR genotyping from short-read data. To some extent, this ambiguity is intrinsic to short reads, because closely related alleles may not always be separated by distinguishing variants within the sequenced fragments. In principle, some ambiguities could be resolved by borrowing information from linked intron variants. In practice, however, this strategy was of limited value because intronic regions often contained substantial sequence variation not represented in the reference database, resulting in a high density of mismatches that made reliable phasing difficult. This problem was further compounded by the fact that many alleles in the database lack complete intronic sequence information. Consequently, adding intronic sequence alone is unlikely to fully resolve these ambiguous genotypes. A more promising strategy may be to incorporate cohort-level information for statistical phasing, as implemented in PING, which we plan to explore in future work.

Overall, KIR*BLOOM achieved strong performance across the 97 real-world WGS samples analyzed in this study. The main remaining limitation appears to be incomplete representation of KIR diversity in current reference resources rather than the inference framework itself. Continued expansion of the IPD-KIR database, together with advances in long-read sequencing, should enable more complete characterization of full-length KIR haplotypes, including intronic and structurally complex alleles, across larger and more diverse cohorts. As long-read sequencing becomes increasingly scalable and cost-effective, these data should substantially improve reference completeness and reduce many of the ambiguities that still limit high-resolution genotyping.

## Methods

### KIR genotyping model

Our method performs KIR genotyping from short-read sequencing data based on a likelihood model that evaluates read depth and sequencing error on KIR alleles under alternative KIR genotype configurations. As a first step, KIR*BLOOM identifies relevant read pairs, maps them against the KIR*BLOOM allele database, and heuristically reduces the solution space by identifying and removing KIR alleles that are likely absent from the genotyped sample. Within the reduced solution space, KIR*BLOOM determines the likely copy number of each gene, followed by selection of actual KIR alleles, in a manner bounded by the inferred copy numbers. In the following, we first describe the core statistical model of KIR*BLOOM that is used for copy number inference and allele selection; the heuristic steps for reducing the solution space and the method for identifying KIR-related read pairs are described later.

Let R denote the set of KIR read pairs, let *B* = {*b*_1_, …, *b*_*n*_} denote the set of KIR alleles from the KIR*BLOOM reference database remaining after heuristic filtering, and let *c* be a copy number vector of length *n*, where the *i*-th element represents the copy number of KIR allele *b_i_*, defined to be between 0 and 4. Our aim is to find a configuration of *c* that maximizes the likelihood of the coverage and read error profiles of the alleles in *B*, where we account for alignment ambiguity of the read pairs in R by considering the probabilities of their alignment locations conditional on *c*.

In the following, we first describe the initial KIR read mapping step and how we probabilistically distribute individual read pairs r over the alleles in B, conditional on c. We then describe how we aggregate read alignments at the level of the KIR alleles in B, obtaining a coverage and read error profile for each allele in B, conditional on c.. Last, we describe how assess consistency between the KIR alleles’ coverage and read error profiles that result from a particular instantiation of c on the one hand and sample genome-wide sequencing depth and sequencing error rate on the other hand using a likelihood model, and how we explore the space of possible configurations of *c* to maximize this likelihood.

### Computation of the alignment probability distribution (*A_r_*) for an individual read pair r

Let 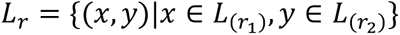 be the set of possible paired-end mapping locations of read *r* against the set *B* of KIR alleles remaining after heuristic filtering; *L*_*r*_ is generated by mapping KIR reads against the KIR*BLOOM database in single-ended mode, considering all possible pairs of individual read alignments, and by filtering these for correct strandedness, insert size etc.; for details. see section “Alignment of KIR-related read pairs” for details. The read likelihood P(r|l), representing the probability of observing *r* if it originated from location *l*∈*L_r_* on KIR allele *b*(*l*) is defined as:

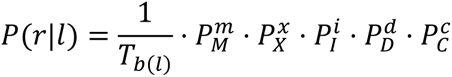

where *m*, *x*, *i*, *d*, *c* represent the number of match, mismatch, insertion, deletion and clipping operations in alignment *l*, respectively, and *P_M_*, *P_X_*, *P_I_*, *P_D_*, *P_C_* denote the probabilities of observing a match, mismatch, insertion, deletion, and clipping event, respectively (see section “Parameter estimation”). *T_b(l)_* is the length of allele *b*, included as a normalization term to account for differences in allele lengths.

We then define the probability that read pair *r* emanated from *l* as:

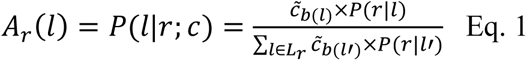

with 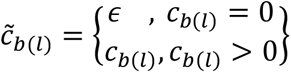

Setting *ɛ* to a small value (> 0) allows for the assignment of reads to alleles with copy number 0. *A*_r_(*l*) for alignments on alleles *b*(*l*) with copy number 0 will always be small, unless *r* has no other possible alignment locations. Reads aligning to alleles with copy number 0 are considered inconsistent with the current configuration of *c* and therefore penalized during later stages of inference. The specific value of *ɛ* varies during different stages of inference (see below).

### Computation of allele alignment profiles P_b_

Denote the allele alignment profile for allele *b* as 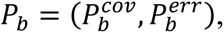 where 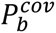 and 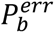 are two vectors of length *T_b_* that represent the coverage and sequencing error for all positions on *b*. Formally, for position *i* of allele *b*, let *L_b,i_* denote the set of alignment locations of all reads that overlap position *i*. Each *l* ∈ *L*_*b*,*i*_ has an associated read pair *r*(*l*), and the probability *l* is the alignment location of read pair *r*(*l*) is *A*_*r*(*l*)_(*l*) (see previous section).

Coverage at positions *i* is defined as

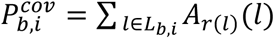

That is, 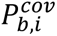 is the expected value of reads covering position i, where the expectation is taken over A_r_ for all relevant reads. Similarly, cumulative read error is computed as 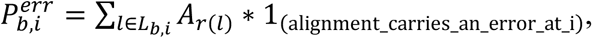 where 1_(_alignment_carries_an_error_at_i) is an indicator function that is 1 if and only if the aligment l carries an error at position i.

### Likelihood function for scoring allele alignment profiles

KIR*BLOOM uses a likelihood model to assess consistency between the alleles’ coverage and read error profiles, conditional on c, and genome-wide estimates of sequencing depth (adjusted for copy number) and sequencing error.

Define *P*_*B*_ = {*P*_*b*_: *b* ∈ *B*} as the set of allele alignment profiles of the alleles in *B*. The likelihood of these, computed conditional on *c*, is defined as

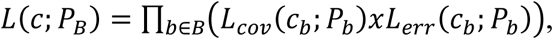

where *L*_*cov*_(*c*_*b*_; *P*_*b*_) and *L*_*err*_(*c*_*b*_; *P*_*b*_) represent coverage and read error likelihoods, respectively; the specific functional form of the utilized likelihood depends on the stage of inference (copy number inference or allele selection; for details, see section “KIR genotype inference”).

The coverage likelihood *L_cov_*(*c_b_; P_b_*) for an allele *b* is computed in a window-based manner. Windows are defined separately for intronic and exonic positions and ignoring the UTR; for full details, see section “Window construction”. Let *w* denote a window, defined as a set of (exclusively intronic or exonic) positions on allele *b*, and let *W*_b_ be the set of all windows associated with *b*. By definition, the union of the positions in *W*_b_ coveres the non-UTR positions of *b*; UTRs sequences are ignored because they are generally less well-represented in IPD-KIR. Let is_exon(*w*) be an indicator function that equals 1 if and only if *w* is an exonic window, and 0 otherwise. The average coverage of a window *w* is defined as

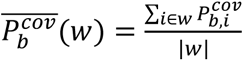

The coverage likelihood *L_cov_*(*c_b_; P_b_*) is defined as

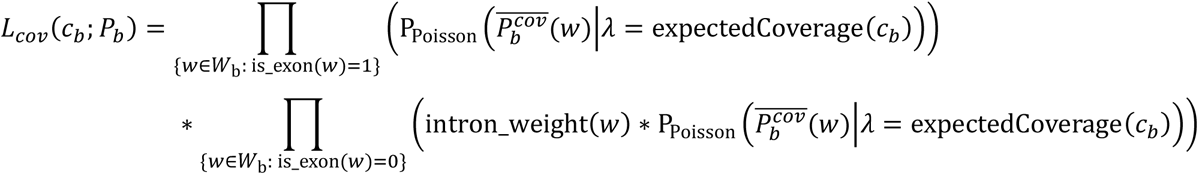

That is, *L*_*cov*_(*c*_*b*_; *P*_*b*_) is computed over the windows of *b*, treating their average coverages as independent, and separately for intronic and exonic windows. The assumed expected coverage for each window depends on the underlying allele’s copy number *c*_*b*_; if *c*_*b*_ > 0, expectedCoverage(*c*_*b*_) is defined as *c*_*b*_ ∗ *D*, where *D* is the estimated haploid genome-wide sequencing depth (see section “Parameter estimation”); if *c*_*b*_ = 0, expectedCoverage(*c*_*b*_) is defined as 0.01 ∗ *D*, penalizing read alignments to alleles with copy number 0 while always retaining a non-zero likelihood for such alleles. Finally, intron_weight(*w*) determines whether the inference process uses coverage information from intronic windows; it is defined as 0 or 1, depending on the stage of inference (section “KIR genotype inference”).

For alleles with c_b_ > 0, the error likelihood *L_err_*(*c_b_; P_b_*) for an allele *b* is computed over the exonic positions of the allele. Let *E_b_* denote the set of exonic positions, and define

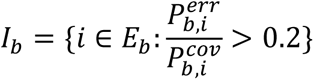

that is, *I_b_* represent the set of exonic positions with error rate > 0.2. We then define *L_err_*(*c_b_; P_b_*) as

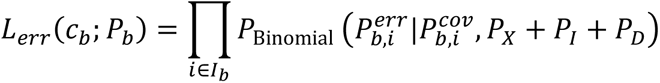

where *P_X_*, *P_I_*, and *P_D_* denote the empirical probabilities of observing a mismatch, insertion, and deletion event, respectively (see section “Parameter estimation”). For alleles with c_b_ = 0, *L_err_*(*c_b_; P_b_*) is defined as 1. Of note, our error model implicitly assigns a likelihood of 1 to all exonic positions with an error rate ≤0.2; this is motivated by increasing robustness against background sequencing error and also to reduce the overall number of positions considered for alleles with c_b_ > 0, as each considered position implicitly penalizes the presence of the allele.

### KIR genotype inference

KIR genotype inference is carried out in two stages, copy number inference and 5-digit allele selection.

- During copy number inference, sequencing errors are ignored (*L*_*err*_(*c*_*b*_; *P*_*b*_) ≔ 1), intronic positions contribute to *L*_*cov*_ (i.e. *intron*_*weight*(*w*) = 1), and *ϵ* = 10^−4^. To optimize *L*(*c*; *P*_*B*_) over the space of possible *c*, we employ a forward selection strategy. We initialize *c*^0^ as a vector of length |*B*|, with 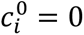 for all *i* ∈ {1. . |*B*|}. During iteration *k*, we then evaluate, for all *i* ∈ {1. . |*B*|}, *L*(*d*^*i*^; *P*_*B*_), where *d*^*i*^ = *c*^*k*−1^, except for 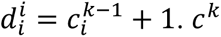 is then set to the *d*^*i*^that achieved the highest likelihood. The process terminates when the maximum log likelihood of iteration *k* does not increase compared to iteration *k* − 1. To accelerate inference, all copy number inference steps are executed on a reduced subset of alleles *B*′ ⊆ *B*, selected based on an EM algorithm that estimates an overall abundance *F*_*b*_ for each allele *b* ∈ *B* in the set of all reads *R*. *B*′ is constructed from *B* in a gene-wise manner, retaining, from all alleles *B*_*g*_ belonging to KIR gene *g*, the allele 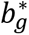 that achieved the highest abundance 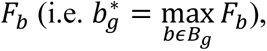 as well as all other alleles 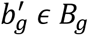 with 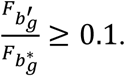
- During 5-digit allele selection, sequencing errors are taken into account (using the definition of *L*_*err*_(*c*_*b*_; *P*_*b*_) given in the previous section), introns are not used in the process of computing *L*_*cov*_ (i.e. *intron*_*weight*(*w*) = 0), and *ϵ* = 10^−300^. Like copy number inference, allele selection operates on a reduced subset of alleles *B*′′ ⊆ *B*; *B*′′ is constructed from *B* by removing all elements corresponding to genes with estimated copy number 0. Let *c*^AS^denote the copy number vector of length |*B*^′′^| used for allele selection; we initialize *c*^AS^ by setting, for all genes with estimated copy number 1, the element corresponding to the allele with the highest EM-estimated abundance to 1; for genes with estimated copy number > 1, the elements corresponding to the alleles selected during copy number inference are set to the corresponding estimated copy number; and all other elements of *c*^AS^ are set to 0. To optimize *L*(*c*^AS^; *P*_*B*′′_) over the space of possible *c*^AS^, we employ a multi-level allele swapping strategy; copy number per gene is held constant over all steps. First, we determine the set of genes *G*^∗^for which alleles currently present in *c*^AS^may not represent the true genotype. To do so, we check, for all alleles *b* belonging to *g* (i.e. for all *b* ∈ *B*^′′^_*g*_) whether i) there is at least one exonic position with an error rate ≥ 0.2 (i.e. whether |*I*_*b*_| > 0) or whether ii) there is at least one window *w* ∈ *W*_*b*_ for which 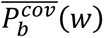 deviates significantly (two-sided Poisson test, *p* = 0.01) from expectedCoverage 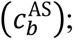 if any of the two conditions is met for alleles *b*, the corresponding gene *g* is added to *G*^∗^. Second, *L*(*c*^AS^; *P*_*B*′′_) is optimized using one-allele swapping between alleles belonging to different 5-digit allele groups. Swapping is implemented in a gene-wise manner, i.e. during each iteration, we sequentially process all *g* ∈ *G*^∗^. Within each gene *g*, we process all pairs (*b*^1^, *b*^2^) of alleles in g (i.e. (*b*^1^, *b*^2^) ∈ {*B*^′′^_*g*_}^2^) for which *b*^1^ ≠ *b*^2^, for which *b*^1^ has a current copy number > 0, and for which *b*^1^ and *b*^2^ belong to different 5-digit allele groups; we then decrease the copy number of *b*^1^ by 1 and increase the copy number of *b*^2^ by 1 (i.e., we swap a copy of *b*^1^ for a copy of *b*^2^) and evaluate the likelihood. After having processed all genes, we change *c*^AS^to the configuration that achieved the highest likelihood (or leave *c*^AS^ unmodified, if no swapping operation was associated with an increase in likelihood). One-allele swapping terminates if one complete iteration over all *g* ∈ *G*^∗^ did not result in any swaps. Third, *L*(*c*^AS^; *P*_*B*_^′′^) is optimized using two-allele swapping. Like one-allele swapping, two-allele swapping is implemented in a gene-wise manner and only between alleles belonging to different 5-digit allele groups. Within each gene *g*, we process all pair of pairs ((*b*^1^, *b*^2^), (*b*^3^, *b*^4^)) of alleles in g (i.e. (*b*^1^, *b*^2^) ∈ {*B*^′′^*_g_*}, (*b*^3^, *b*^4^) ∈ {*B*^′′^*_g_*}^2^) for which *b*^1^ ≠ *b*^2^; *b*^3^ ≠ *b*^4^; *b*^1^ and *b*^2^ have a current copy number > 0; *b*^3^ and *b*^4^ have a current copy number of 0; and for which the sets of 5-digit allele groups associated with (*b*^1^, *b*^2^) and (*b*^3^, *b*^4^) are not identical. For each pair of pair, we decrease the copy numbers of *b*^1^ and *b*^2^by 1, increase the copy numbers of *b*^2^and *b*^3^by 1, and evaluate the likelihood. After having processed all genes, we change *c*^AS^ to the configuration that achieved the highest likelihood (or leave *c*^AS^ unmodified, if no swapping operation was associated with an increase in likelihood). Two-allele swapping terminates if one complete iteration over all *g* ∈ *G*^∗^ did not result in any swaps.

### EM algorithm

We use the EM algorithm to reduce the size of the allele set considered for copy number inference, and also for seeing the allele selection step. Briefly, each KIR allele *b* ∈ *B* is assumed to have an abundance *F*_*b*_ over all read pairs *r* ∈ *R*, and each read pair *r* is assumed to have a hidden variable that indicates the KIR allele that read pair *r* emanates from. We define *P*(*r*|*b*) as identical to the highest-likelihood mapping location on allele *b*, i.e. 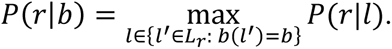 The probability that read *r* emanated from allele *b* is then defined as 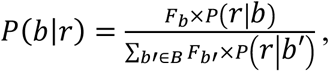 and the overall likelihood is *L*(*R*) ≔ ∏_*r*∈*R*_ *F*_*b*_ × *P*(*r*|*b*) *F*_*b*_ is iteratively maximized using EM, i.e. by computing *P*(*b*|*r*) based on the current estimate of *F*, followed by updating *F* based on *P*(*b*|*r*) for all *r* ∈ *R*, until the increase in log *L*(*R*) falls below 0.01.

### Parameter estimation

A copy-number–stable background region of 2Mbp on GRCh38 chromosome 17 (specific coordinates: chr17:74Mb-76Mb) (Prodanov et al. 2025) is used for parameter estimation at multiple stages of the pipeline. Alignment event probabilities are estimated by counting matches (m), mismatches (x), insertions (i), deletions (d), and clipped bases (c) from primary paired-end read alignments and by normalizing these counts by their total to obtain *P_M_*, *P_X_*, *P_I_*, *P_D_*, *P_C_*, respectively. The chromosome 17 control region is also used for estimating haploid genome-wide sequencing depth (parameter *D*) as average per-base sequencing depth, divided by 2.

### Identification of KIR-related read pairs

KIR*BLOOM supports BAM and CRAM files aligned to 1000 Genomes GRCh38 reference genome, as well as raw reads in FASTQ format; if FASTQ format is used, reads are first aligned to the 1000 Genomes reference using bwa-mem2.

Based on GRCh38 read alignments, a first set of candidate KIR read pairs is obtained by extracting all pairs of reads for which at least one member read aligns into the KIR region on chromosome 19 (coordinates) or onto the alternative LRC/KIR haplotypes, as well as read pairs with at least one unaligned member read.

The candidate KIR read set is further refined by aligning, using minimap2, the extracted pairs against a combined reference genome that comprises i) the B38 reference with all KIR gene locations masked; ii) the T2T reference genome with all KIR gene locations masked; iii) the KIR*BLOOM allele database. Read pairs with properly paired primary alignments to the KIR*BLOOM reference database represent high-confidence KIR reads; these therefore constitute the final set of read pairs used for KIR genotyping. Masking of the two reference genomes was carried out based on manually curated KIR gene annotations (see Section “Annotation of KIR loci in GRCh38 and T2T references”). Combining the T2T and GRCh38 references during the refinement stage reduces nonspecific mappings to repetitive elements that would otherwise be propagated into the KIR genotyping step.

### Alignment of KIR-related read pairs

For each read pair *r* = (*r_1_*, *r_2_*)ns sets 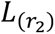 and ￼ for *r_1_* and *r_2_* by mapping them in single-ended mode against the KIR*BLOOM reference database, using minimap2 in short-read mode (-ax sr --secondary-seq --MD --eqx The set of possible paired-end alignment locations is then defined as ￼, where only members that satisfy the following constraints are retained: (i) both alignments map to the same KIR allele, (ii) the alignments have inward-facing orientations on opposite strands, and (iii) the inferred insert size lies within the interval [*μ*−4*σ*, *μ*+4*σ*], where *μ* and *σ* the considered set of alignment locations for read ￼generated by filtering *B* to KIR alleles present in the reduced allele set *B*.

### Heuristic reduction of the KIR allele solution space

After identifying KIR-related reads, the first step of KIR*BLOOM is to heuristically reduce the search space by identifying KIR alleles that are likely not present in the genotyped sample, based on a two-step filtering strategy. The KIR alleles that survive the heuristic filters become the allele set *B* that goes into the main genotyping model.

During the first filtering step, we remove alleles *b* that have at least one “core” position with alignment depth 0 when considering the set of all possible KIR read pair alignment locations 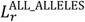 for all reads *r* ∈ *R*. The “core” positions of an allele are defined as positions that lie i) outside of the UTR region of the allele and b) outside of the first or last 15% of the allele’s sequence; note that the “alignment depth” criterion used here includes non-match alignment operations.

During the second filtering step, we focus on positions that discriminate between KIR alleles of the same gene, and we rank (and then remove) alleles according to their read support at such positions. Formally, for gene *g*, let the set *Z* denote the columns in the MSA of the alleles of *g* that discriminate between alleles (i.e., which contain more than one base) and which are consistently annotated with respect to the intron/exon status of the included alleles at this position in the MSA. To score allele *b* of gene *g*, we iterate through the set *Z*, removing all columns at which the sequence of *b* that was interpolated (during the construction of the KIR*BLOOM reference database; see section “Construction of the KIR*BLOOM allele reference database”); at which *b* carries a “gap” character; or which are part of the UTR. Let the set *Z*^*b*^ denote the columns in the MSA that remain after filtering and that are used for scoring allele *b*; furthermore, let 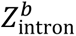 and 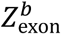 denote the intronic and exonic subsets of *Z*^*b*^, respectively. Alleles with 0 remaining positions for evaluation (i.e. alleles with |*Z*^*b*^| = 0) are always retained. For the other alleles, we count, at each position *z* ∈ *Z*^*b*^, the number of “match” characters across all possible read alignments at the considered position (i.e., based on the set 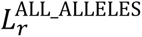 for all reads *r* ∈ *R* that overlap the position), and carry out a statistical hypothesis test against the null hypothesis that the observed number of matches can be explained by sequencing error alone (one-sided Poisson with *λ* = 1 and significance threshold depending on the relative location of the position within *b*, see below). Positions at which the null hypothesis is rejected are counted as “supported”, and positions at which the null hypothesis is not rejected are counted as “not supported”. For each allele, we separately compute the total number of unsupported exonic and intronic positions. If there is a single allele that has a lower count of unsupported exonic positions than all other alleles within the set of considered alleles with |*Z*^*b*^| > 1, this allele is retained, and all other alleles with at |*Z*^*b*^| > 1 are removed. Otherwise, we consider the set of alleles that achieve the same minimum number of unsupported exonic positions and retain i) all alleles with 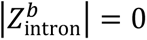 and ii) all alleles with 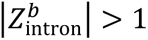 that have the minimum number of unsupported intron positions. The significance threshold used for the Poisson test is *p<0.05* for all positions that fall into the central 70% region of each allele, and p<0.1 otherwise.

### Annotation of KIR loci in the GRCh38 and T2T references

We annotated KIR loci in both the GRCh38 and T2T reference genomes based on available reference annotations (RefSeq for GRCh38 and JHU RefSeq v110 for T2T), Immuannot, and KIR allele mappings generated by aligning the IPD-KIR database with interpolated introns and UTRs (see Section “Construction of the KIR-BLOOM allele reference database”) to the reference genomes. For each reference genome, we took the union of KIR loci identified by these three sources. For each locus, gene labels were determined based on Immuannot annotations, while locus start and end positions were defined using the KIR allele mappings. In addition, the annotation results for each locus were manually curated.

### Construction of the KIR*BLOOM allele reference database

The KIR*BLOOM allele database was constructed using the IPD-KIR database (December 2025 release) together with novel alleles identified by Immuannot from the T2T and GRCh38 genomes (see Section “Annotation of KIR loci in the GRCh38 and T2T references”). The IPD-KIR database contains 2,286 alleles available only as coding sequences (CDS); for these, intronic sequences were interpolated from the most closely related intron-and-exon-complete alleles. Finally, UTR sequences for all alleles were completed, based on either i) the UTR sequence of the allele that intron interpolation was based on, if this allele contained a UTR and if the allele with incomplete UTR was originally only available as CDS; or ii) based on inserting UTR sequence from an arbitrary UTR-complete allele. Details of the KIR*BLOOM allele reference construction are provided in the Supplementary Notes.

### Window construction

The coverage component *L*_*cov*_(*c*_*b*_; *P*_*b*_) of the KIR*BLOOM likelihood is computed in a window-based manner rather than at single-base resolution to reduce sensitivity to local read coverage and to account for statistical non-independence between neighboring positions. Windowing is carried out separately for each gene *g* and, for each allele, exonic and intronic regions are partitioned separately, and UTR regions are excluded. The exonic portion of each allele *b* ∈ *B*_*g*_ is partitioned into contiguous windows of length 150bp for all windows except for the last window; if the last window has a size of less than 100bp, it is combined with the previous window. If, after having partitioned all alleles of gene *g*, there is a difference between the alleles of *g* with respect to the number of windows, windowing is repeated with a fixed target number of windows per allele, equal to the minimum window count achieved during the first iteration. During this second iteration, once the number of 150bp windows has achieved its allowed maximum, the last windows is extended until all relevant positions are included. Intronic positions are partitioned analogously.

### Variants correction

To refine the inferred allele sequences, we applied a variant-correction procedure based on read support at exonic sites. First, for each inferred allele, we calculated the fraction of non-reference observations at each exonic position as the sum of mismatches, insertions, and deletions divided by the total depth at that site. Positions with an error fraction greater than 0.75 were flagged as candidate correction sites, and the most strongly supported alternative allele in the read pileup was recorded. Second, for genes with two inferred alleles, we used the gene-specific multiple sequence alignment to identify ungapped exonic positions at which the two alleles differed. At each such site, we compared the read support for the two allele-specific bases. When more than 85% of the support favored one base, the alternate allele was corrected to match the supported base, effectively collapsing unsupported heterozygous differences into a homozygous call. All candidate corrections were written as VCF records for downstream sequence updating.

### Development and validation data

Whole-genome sequencing data were obtained from two publicly available sources. First, pre-aligned CRAM files were downloaded from the European Nucleotide Archive (ENA) FTP repository. These datasets correspond to samples for which HGSVC3 assemblies are available. All such CRAM files were aligned against the GRCh38 full analysis set plus decoy and HLA sequences reference genome, which was obtained from https://42basepairs.com/download/s3/1000genomes/technical/reference/GRCh38_reference_genome/GRCh38_full_analysis_set_plus_decoy_hla.fa. Second, additional whole-genome sequencing datasets were obtained from samples generated by the Human Pangenome Reference Consortium (HPRC). For these samples, paired-end whole-genome sequencing data (fastq files) were downloaded individually from the NCBI by querying the Sequence Read Archive (SRA) for available WGS paired-end sequencing runs.

For samples available as CRAM files, reads had already been aligned using bwa-mem2 (v2.2.1); therefore, reads mapping to KIR-associated regions were directly extracted from these alignments. For the HPRC datasets, raw sequencing reads were first aligned using bwa to GRCh38 full analysis set reference genome, including decoy and HLA sequences, after which KIR-related reads were similarly extracted. The primary KIR locus on chromosome 19 (chr19:54,025,634–55,084,318) was selected and extended by ±1 kb to capture flanking regions. In addition, alternative reference sequences corresponding to the leukocyte receptor complex (LRC) were obtained from Genome Reference Consortium resources (https://www.ncbi.nlm.nih.gov/grc/human) by selecting alternative contigs whose names contained the string LRC-KIR. Finally, a non-KIR region on chromosome 17 (chr17:74,000,000–76,000,000) was included as a background reference region for downstream analyses (Prodanov et al. 2025).

### KIR read identification

To identify reads for KIR genotyping, we constructed a joint reference genome consisting of the T2T and GRCh38 reference genomes, in which KIR loci were masked, together with the KIR*BLOOM allele database. Input reads were mapped against this joint reference genome using BWA in paired-end mode. Read pairs for which both reads had a primary alignment to the KIR-BLOOM reference database were retained for downstream KIR genotyping. Combining the T2T and GRCh38 references at this stage reduces nonspecific mappings to repetitive elements that would otherwise be propagated into the KIR genotyping step.

### Constructing the ground truth

Ground-truth KIR genotypes were derived from haplotype-resolved genome assemblies available for each sample using Immuannot and IPD-KIR release 2.15.0. After running Immuannot with default parameters on the two phased haplotypes of each sample, we parsed the resulting maternal and paternal GTF files to extract the consensus CDS annotation for each KIR gene. For every annotated KIR copy, we recorded the consensus CDS sequence together with its reported CDS distance, and then combined the maternal and paternal annotations to construct the ground-truth CDS set for each sample. From these annotations, we also derived the ground-truth gene copy number by counting the total number of annotated KIR copies across the two haplotypes. These CDS-resolved and copy-number truth sets were then used as the reference for downstream performance evaluation.

### Performance evaluation

Performance was evaluated separately at the gene copy-number level and at the allele level for each sample and gene. For copy-number evaluation, the predicted and true copy numbers were compared directly. The number of correctly inferred copies was counted as true positives, defined as the minimum of the true and predicted copy number for a given gene. Any excess predicted copies were counted as false positives, and any missing copies were counted as false negatives. An exact copy-number match was recorded when the predicted copy number was identical to the true copy number. At the allele level, after variant calling, allele-level performance was assessed using the CDS sequence of allele calls from while preserving allele multiplicity. For each gene, correctly predicted CDS copies were counted as true positives, missing true CDS copies as false negatives, and additional predicted CDS copies as false positives. This ensured that partial recovery of a duplicated CDS was scored proportionally; for example, if a CDS was present at copy number two in the truth set but predicted at copy number one, this contributed one true positive and one false negative. Exact allele match was defined as complete agreement in both CDS identity and copy number.

For both copy-number and allele-level evaluations, precision was calculated as TP / (TP + FP), recall as TP / (TP + FN), and the Jaccard index as TP / (TP + FP + FN). All metrics were reported as percentages. Exact-match performance was calculated as the proportion of evaluated sample–gene combinations for which the prediction matched the truth exactly.

### Benchmark methods

T1K v1.0.9-r239 was benchmarked using the IPD-KIR 2.15 database. To harmonize allele resolution with the evaluation framework, T1K was run with --alleleDigitUnits 5, producing 5-digit genotype calls in genotype.tsv. These calls were subsequently refined with t1k_copynumer.py script, and the resulting copy-number-corrected 5-digit alleles were used for evaluation before variant correction. For evaluation after variant correction, the T1K allele.tsv file and VCF output were used to reconstruct 7-digit representative alleles. Briefly, the corrected 5-digit calls were mapped to the corresponding 7-digit representative alleles reported by T1K, after which sequence variants from the VCF were applied. For alleles with copy number greater than one, variants were assigned to ceil(Freq × copy number) copies. When multiple variants were present, this assignment was performed iteratively for each variant, with copies selected arbitrarily because haplotype-specific phase information was not available. GraphKIR was run using the supplied index (IPD-KIR 2.10). Geny was also run with the available database (IPD-KIR 2.12). In both cases, the resulting allele and copy-number predictions were used for downstream benchmarking after converting reported allele names to their corresponding IPD-KIR 2.15 nomenclature using Allelelist_history.txt from IPD-KIR.

## Availability of data and materials

KIR*BLOOM source code is available in https://github.com/YomnaGohar/KIR-BLOOM Construction of KIR*BLOOM allele database is available at https://github.com/YomnaGohar/Interpolating-Complete-Sequences-for-KIR-Alleles

## Authors’ contributions

Y.G. and A.D. developed the KIR*BLOOM method and wrote the manuscript. Y.G. implemented the method and performed the analyses. A.D. supervised the study. A.D.G. contributed at an early stage of the project by helping to identify the optimal modeling strategy and define the evaluation metrics. K.M.K. performed the PING benchmarking analyses. P.J.N. contributed through scientific discussion and advice. All authors reviewed and approved the final manuscript.

