## Supplementary Figures for "KIR*BLOOM: Accurate KIR genotyping using a new copy number-aware integrated genotype likelihood framework"

### Supplementary Figure 1

Overview of KIR*BLOOM pipeline

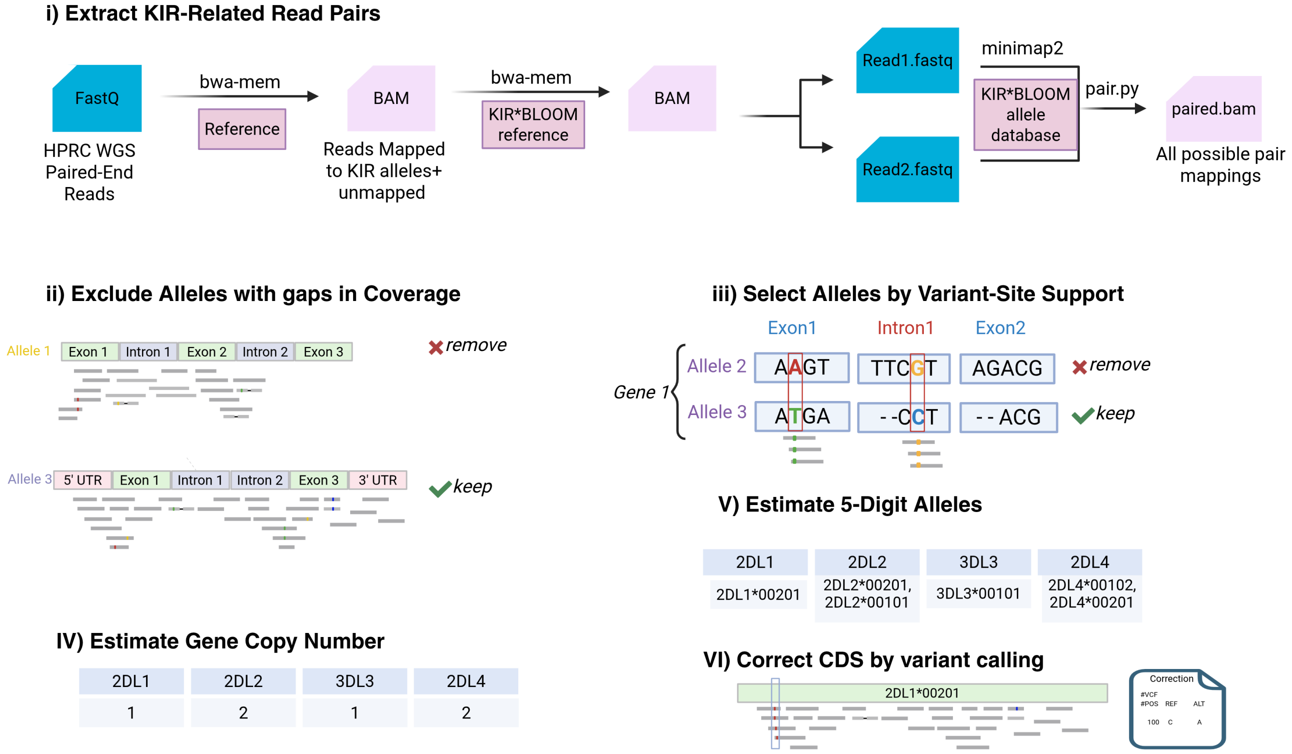


### Supplementary Figure 2

Evaluation of gene copy-number inference performance.

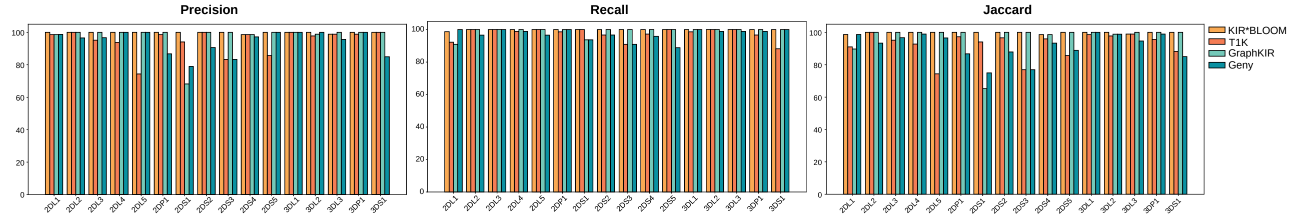

**Supplementary Figure 3**

Schematic representation of an ambiguous genotype (top). Coverage and error profiles for the alleles predicted by KIR*BLOOM (left) and the true genotype (right) for KIR3DL3.

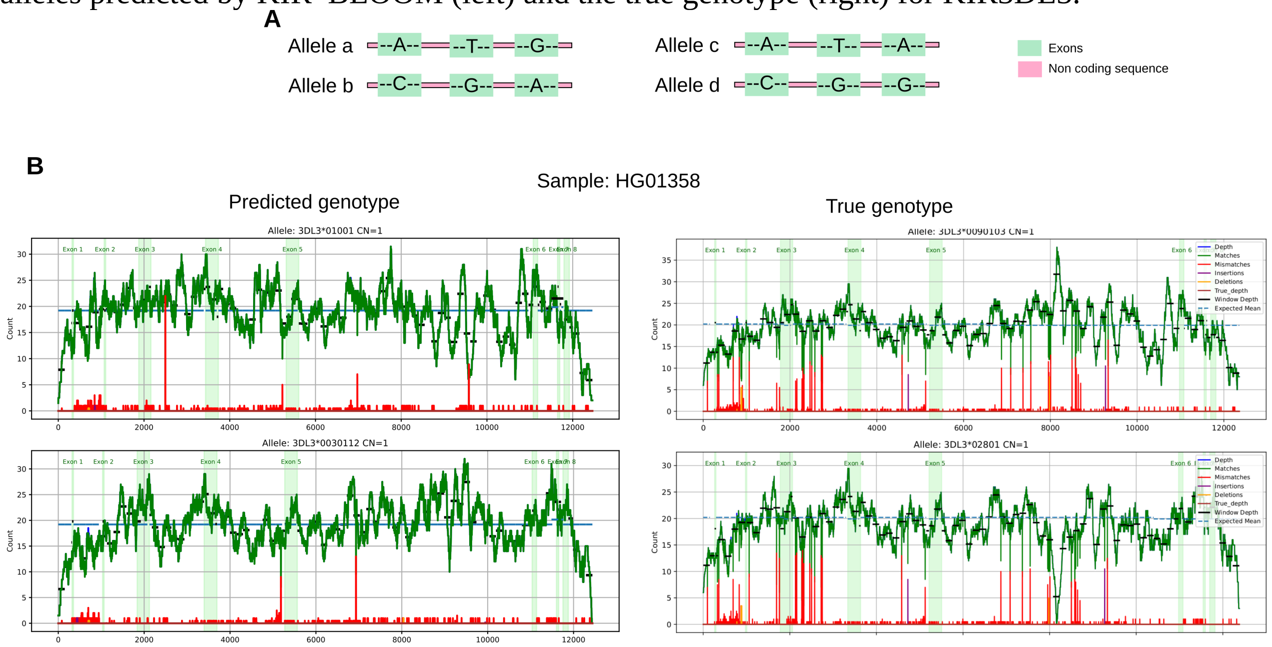
