## Supplementary Notes for "KIR*BLOOM: Accurate KIR genotyping using a new copy number-aware integrated genotype likelihood framework"

### Reference Construction

For accurate KIR genotyping, KIR-related reads must first be extracted, which requires the use of a reference genome. In this study, the reference consisted of two components: (1) a human reference genome in which KIR loci were masked, and (2) a database of curated KIR allele sequences.

For component (2), we downloaded the IPD-KIR database (December 2025 release), which contains both complete and partial allele sequences, including entries represented only by coding DNA sequences (CDS). The first step involved constructing full-length alleles by interpolating missing intronic and untranslated regions (UTRs) for CDS-only entries, as described below.

For component (1), we annotated KIR loci in both the GRCh38 and T2T (CHM13) human genome assemblies using three complementary approaches. First, curated public annotations were employed: RefSeq annotations for GRCh38 and the JHU RefSeq v110 annotations generated with Liftoff v5.2 for T2T. Second, we applied the **Immuannot** tool to both assemblies to obtain KIR-specific feature predictions. Third, all KIR alleles from the IPD-KIR database were mapped to both assemblies to delineate KIR locus boundaries precisely.

All candidate KIR loci were manually inspected to resolve discrepancies between annotation sources, such as overlapping or conflicting gene labels (see Supplementary File X for examples). When inconsistencies arose, Immuannot annotations were prioritized based on their concordance with allele mappings. For each confirmed locus, masking coordinates were defined as the minimum start and maximum end positions across all aligned alleles, thereby ensuring inclusion of variable UTR regions and preventing residual KIR-related sequences from attracting spurious mappings. This process produced masked reference genomes that retained intergenic and non-KIR regions while comprehensively excluding all KIR loci.

Novel KIR alleles identified during annotation were extracted using a custom Python script based on the GTF file generated by Immuannot. These novel alleles, verified through manual inspection and sequence comparison, are provided in **Supplementary Table Novel_alleles.ods**.

Novel alleles identified during the annotation step were incorporated into the corresponding multiple sequence alignment (MSA) files for each KIR gene. Each MSA contained the full set of alleles from the IPD-KIR database, including those with interpolated intronic and untranslated regions (UTRs). Novel alleles were added to the existing MSAs using **MAFFT** with the `--add` and `--reorder` options. Following integration, the 5′ and 3′ UTRs of all alleles within each gene were visually inspected and standardized to ensure uniform UTR lengths, except in cases where extensive structural heterogeneity or numerous alignment gaps exist.

All KIR allele sequences across all genes, together with the GRCh38 and T2T (CHM13) reference assemblies in which KIR loci were masked, were concatenated into a single FASTA file. This comprehensive reference was used throughout the study for read mapping, allele identification, and genotyping analyses.

##### 1. Modify IPD-KIR

The complete sequences of all KIR alleles are not fully known, as some alleles are represented only by their cDNA sequences, which include exons but lack introns, untranslated regions (UTRs), and, in some cases, even terminal exons are partially or completely missing. To achieve a comprehensive set of KIR alleles enabling genotyping at a seven-digit resolution, we interpolated the missing sequences from closely related alleles with complete sequences. To accomplish this, we utilized MSF files from IPD-KIR (December 2024). These MSF files contain multiple sequence alignments for alleles of each KIR gene. There are three types of MSF files: protein MSF, genomic MSF, and nucleotide MSF. The genomic MSF contains the multiple sequence alignments of alleles with complete sequences, except for a few cases (e.g., KIR3DL3*005). The nucleotide MSF contains the multiple sequence alignments of the exons of all known alleles of that gene. The aim is to produce a modified genomic MSF that includes both the complete alleles and the exons of the incomplete alleles, with introns and UTRs (if present) interpolated from closely related alleles, as illustrated in Figure 1S. This step is crucial for enabling the alignment of sequencing reads from whole-genome sequencing (WGS) experiments.

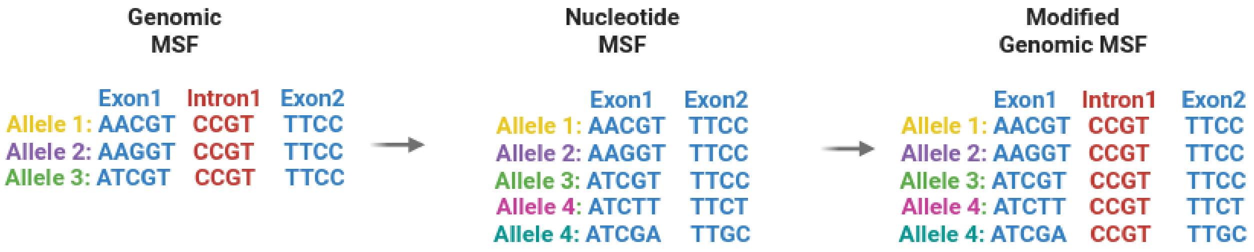


This process was achieved through the following steps:

1. **Transforming Annotations to Account for Gaps in the MSA**:
   We transformed the annotations from the XML file, which contains the coordinates of features, into multiple sequence alignment (MSA) annotations to account for gaps introduced during alignment. This transformation ,using the following function, ensures that the annotations accurately reflect the positions within the MSA.

```
function get_msa_position(msa_sequence, original_position):
  original_idx = 0
  msa_idx = 0
  while original_idx < original_position:
    if msa_sequence[msa_idx] is not '-':
      original_idx += 1
    msa_idx += 1
  while msa_idx < length(msa_sequence) and msa_sequence[msa_idx] is '-':
    msa_idx += 1
  return msa_idx
```

   An example of this process is depicted in Figure 2S.

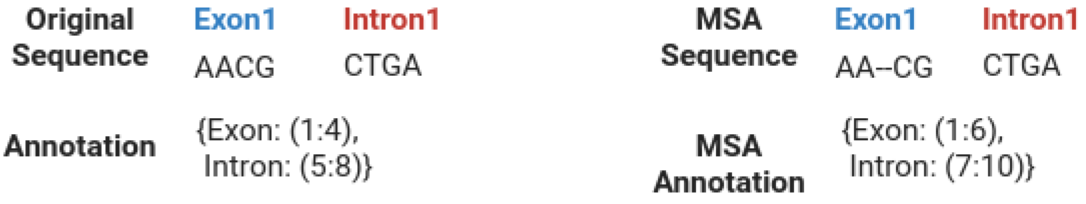


2. **Ensuring No Overlaps Between Features in the MSA Genomic Files**:
   To ensure there are no overlaps between features such as terminal exons and untranslated

regions (UTRs), we standardized the annotations by harmonizing the feature counts and the coordinates of the features across all alleles. For each allele, we ensured the same number of features by adding `None` as placeholders for any missing features at a given position. The feature annotations were then modified: we adjusted the end coordinates of the features to match the maximum end coordinate observed across all alleles at that index. Subsequently, we adjusted the start coordinates so that each feature began one base after the previous feature's end coordinate. These steps are crucial for two reasons: first, to ensure that gaps at the very start or end of features are included in the ranges, and second, to allow the detection of any overlaps between features.

All genes had no overlaps between their alleles' features in the genomic MSA except for KIR2DL3*010 and KIR3DL1*05901 where they had an overlap between last exon and UTR. This overlap was modified using a custom python script. An example of the standardization process is depicted in Figure S3.

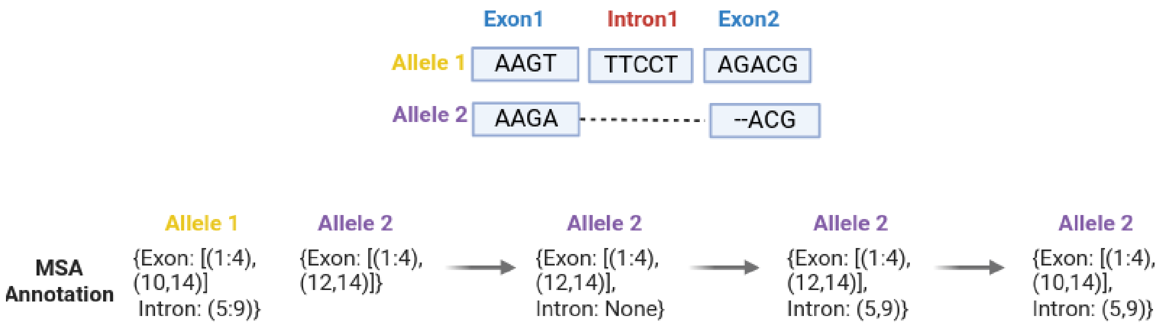


#### Interpolation of Missing Exons in Genomic MSA Using Closely Related Alleles

Some alleles in the genomic msf file contained missing features; however, it is unclear whether this reflects true biological absence or incomplete annotation in the database. Alleles with any identified missing features were therefore not used to complete missing UTR or intronic regions in the nucleotide sequence files. These alleles are reported in the following table.

| Gene | allele | feature/s missing or incomplete |
|---|---|---|
| 2DL5A | KIR2DL5A*00102 | Missing intron(s) |
| | KIR2DL5A*00104 | Missing intron(s) |
| | KIR2DL5A*00105 | Missing intron(s) |
| | KIR2DL5A*0050102 | Missing intron(s) |
| | KIR2DL5A*01201 | Missing intron(s) |
| | KIR2DL5A*01202 | Missing intron(s) |
| 2DL5B | KIR2DL5B*0020102 | Missing intron(s) |
| | KIR2DL5B*0020104 | Missing intron(s) |
| | KIR2DL5B*0020106 | Missing intron(s) |
| | KIR2DL5B*0020201 | Missing intron(s) |
| | KIR2DL5B*003 | Missing intron(s) |
| | KIR2DL5B*00602 | Missing intron(s) |
| | KIR2DL5B*00603 | Missing intron(s) |

| | | |
|---|---|---|
| | KIR2DL5B*0070102 | Missing intron(s) |
| | KIR2DL5B*0080102 | Missing intron(s) |
| | KIR2DL5B*00802 | Missing intron(s) |
| | KIR2DL5B*009 | Missing intron(s) |
| | KIR2DL5B*010 | Missing intron(s) |
| | KIR2DL5B*011 | Missing intron(s) |
| | KIR2DL5B*01301 | Missing intron(s) |
| | KIR2DL5B*01302 | Missing intron(s) |
| | KIR2DL5B*01303 | Missing intron(s) |
| 2DS4 | KIR2DS4*0010103 | Missing intron(s) |
| 2DL4 | KIR2DL4*0010302 | Incomplete exon1 (1 to 28) |
| | KIR2DL4*00104 | Incomplete exon1 (1 to 28) |
| | KIR2DL4*00901 | Incomplete exon1 (1 to 28) |
| | KIR2DL4*0080202 | Incomplete exon1 (1 to 28) |
| | KIR2DL4*0080103 | Incomplete exon1 (1 to 28) |
| | KIR2DL4*010 | Incomplete exon1 (1 to 28) |
| 3DL3 | KIR3DL3*0020201 | Missing exon1 |
| | KIR3DL3*005 | Missing intron(s) |

For alleles that have partially or completely missing exons (which occurs at the terminal sequences), the missing sequences were interpolated from closely related complete alleles of the same gene. We define a complete allele as one that has no missing or partial exons or introns. The following pseudocode describes how a closely related allele is identified:

```
function sequence_difference(seq1, seq2):
    difference = 0
    for each character c1, c2 in seq1 and seq2:
        if c1 ≠ c2:
            difference += 1
    return difference


function find_closest_allele(target_allele, nucleotide_msa_dict,
complete_sequence_annotations):
    target_sequence = nucleotide_msa_dict[target_allele]
    min_difference = infinity
    closest_allele = None

    for each allele, sequence in nucleotide_msa_dict:
        if allele ≠ target_allele:
```

```
        diff = sequence_difference(target_sequence, sequence)
        if diff < min_difference and allele is in complete_sequence_annotations:
            min_difference = diff
            closest_allele = allele
    return closest_allele, min_difference
```

To accurately identify missing or partial exons, we utilized the standardization process described in the previous section. During this process, `None` was added as a placeholder for any missing features (whether exons, introns, or UTRs) across all alleles. This standardization allowed us to systematically detect missing exons, as the presence of `None` indicated that a feature was absent in the sequence. For partial sequences, the identification was based on information derived from the XML file, which includes a `status` attribute for each exon. This attribute specifies whether an exon is complete or partial. However, it is important to note that this `status` attribute is only available for exons; UTRs and introns do not have this attribute in the XML file.

To ensure accurate interpolation of a completely missing exon, we needed to determine whether the missing exon was due to a true biological absence or simply a gap in the data. This determination was also based on the XML file, which contains cDNA sequence data. If the cDNA sequence does not start at position 1, or if there is a gap between the end position of one exon and the start position of the next, it suggests that the absence is due to a gap in the data rather than a biological missingness.

**Interpolation of Missing Introns and UTRs in Incomplete Alleles Using Closely Related Genomic MSAs**

For the alleles in the nucleotide file, which contains the MSA of exons for all alleles—including those in the genomic MSA and those missing introns and UTRs—we interpolated the missing introns and UTRs (if present) from the closest alleles in the modified genomic MSA resulting from the previous section. For each allele, the closest match was found. The interpolation process followed the same steps as those used for interpolating the missing sequences in the genomic MSA.

Briefly, for alleles that are incomplete, we standardized the sequence annotations by adding `None` as a placeholder for the missing exons. We then iterated over the features of the complete sequence of the closest related allele, ordered by their starting positions. If the feature was a UTR or an intron, the sequence was derived from the closely related allele. If the feature was an exon, we checked whether it was missing or partial in the allele. If it was missing, we determined whether the missing exon was due to true biological absence or simply missing data. If the absence was due to missing data, we interpolated the exon from the closely related allele. If the exon was partially missing, we also interpolated the missing part from the closely related allele. If no missingness or partial sequence was present, we inserted the exon as it is.

Importantly, the sequence added during interpolation was the multiple aligned sequence. To ensure consistency, we verified that the multiple sequence alignment of the exons in the closest related allele was identical in both the genomic and nucleotide MSF files before incorporating the modified sequence.
